# A multiscale model of immune surveillance in micrometastases: towards cancer patient digital twins

**DOI:** 10.1101/2023.10.17.562733

**Authors:** Heber L. Rocha, Boris Aguilar, Michael Getz, Ilya Shmulevich, Paul Macklin

## Abstract

Metastasis is the leading cause of death in patients with cancer, driving considerable scientific and clinical interest in immunosurveillance of micrometastases. We investigated this process by creating a multiscale mathematical model to study the interactions between the immune system and the progression of micrometastases in general epithelial tissue. We analyzed the parameter space of the model using high-throughput computing resources to generate over 100,000 virtual patient trajectories. We demonstrated that the model could recapitulate a wide variety of virtual patient trajectories, including uncontrolled growth, partial response, and complete immune response to tumor growth. We classified the virtual patients and identified key patient parameters with the greatest effect on the simulated immunosurveillance. We highlight the lessons derived from this analysis and their impact on the nascent field of cancer patient digital twins (CPDTs). While CPDTs could enable clinicians to systematically dissect the complexity of cancer in each individual patient and inform treatment choices, our work shows that key challenges remain before we can reach this vision. In particular, we show that there remain considerable uncertainties in immune responses, dysfunctional data stratification, and unpredictable personalized treatment. Nonetheless, we also show that in spite of these challenges, patient-specific models suggest strategies to increase control of clinically undetectable micrometastases even without complete parameter certainty.

## 1 Introduction

Metastasis constitutes the most fatal manifestation of cancer and remains among the least understood aspects of cancer pathophysiology. Early metastases are often undetected at primary diagnosis, leading to an increased risk of mortality [1, 2]. Identifying occult metastases in early-stage cancer is crucial for optimal therapy and prognosis [3, 4]. The metastasis process is highly inefficient because individual cancer cells and small clusters arriving in a new tissue must overcome multiple barriers to successfully nucleate a new metastasis [5–7]. At each step of the metastatic cascade, the host immune system can potentially recognize and kill mutant cancer cells, which may also be immunogenic [5, 8]. Nevertheless, cancer cells still find ways to evade host immune surveillance in distant sites. Understanding the nuances of these complex processes could lead to the development of novel therapeutic strategies to benefit patients by targeting otherwise undetected micrometastases to either eliminate them or prevent their further growth.

The interactions between metastatic cancer and the immune system (with or without therapeutic modulation) are inherently multiscale, involving molecular-, cell-, multicellular-, and systems-scale processes at fast and slower time scales, making cancer immunology the quintessential multiscale complex systems problem of our era. Computational models can serve as a “virtual laboratory” to understand and ultimately control the behavior of such complex systems [9, 10]. A Cancer Patient Digital Twin (CPDT), a personalized computational replica of an individual patient’s cancer, leverages these models to simulate disease progression and treatment outcomes. The foundation of CPDTs lies in computational models that facilitate (1) calibrating models with individual patient data, (2) predicting cancer progression across a range of treatment options, and (3) refining models with updated patient measurements over time [11–15]. This paradigm could allow clinicians to systematically assess the progression of an individual patient’s cancer, simulate treatment outcomes, and ultimately choose the best treatment to achieve the patient’s goals. Multiscale models are of special interest for the development of CPDTs, as they can integrate several relevant interactions — often from different temporal and spatial scales — into a unified simulation framework. For instance, multiscale models can simultaneously incorporate molecular and cellular level interactions between cancer cells and the immune system, providing a comprehensive view of the tumor microenvironment. In this work, we develop a multiscale model of metastatic cancer cells growing in epithelial tissue, focusing on the interactions between immune and cancer cells, and use this model to further evaluate key challenges for the nascent field of cancer patient digital twins.

Recently, several mathematical models have been developed to help understand the complex crosstalk between the tumor and the immune system [16–20]. In particular, multiscale hybrid models have been frequently applied due to their modularity and versatility [16, 17, 21–23]. Makarya et al. [24] reviewed how multiscale mechanistic models can represent different aspects of immune system cells and suggest areas of study in immune-cancer interactions. They argue that future efforts in approaches such as multiscale modeling will enable the prediction of the effects of immunotherapy in a range of patientspecific settings and generate new insights into immunotherapy strategies. On the other hand, hybrid multiscale models require estimating parameters that modulate several relevant processes, such as cell-cell communication, gene regulation, cell division, cell death, oxygen consumption, and interactions, resulting in highly complex models. Furthermore, it is challenging to calibrate and validate these models due to the limited availability of patient-specific data (particularly spatially resolved, serial data) and the substantial computational cost of their iterative parameter fitting techniques [25–27]. Consequently, it will be essential to integrate machine learning and high-performance computing (HPC) techniques when creating virtual patient models [25, 28].

In recent work [15, 23, 29], we developed a multiscale agent-based model of innate and adaptive immune cell interactions with COVID-19 infected lung tissue, along with immune cell trafficking between an infected site and the lymphatic system. In this paper, we adapted and extended this model to create a multiscale agent-based model of micrometastases with local and systems-scale immune interactions, including mechanics-based cell death [30–33], secretion of pro-inflammatory cytokines, immune cell recruitment and infiltration.

Using HPC-based simulations and employing an ensemble modeling approach (where multiple simulations are aggregated into statistically summarized results), we verified that the model is capable of capturing clinically salient outcomes including uncontrolled growth, partial tumor control, and complete tumor elimination. Through our virtual experiments, a significant observation emerged: the majority of our patients displayed non-unique outcomes across experimental replicates, underscoring the substantial uncertainty inherent in immune response dynamics. Even in the case of a “thought experiment” where a virtual patient’s parameters are known with complete certainty, the final clinical outcome cannot be determined in advance due to the inherent stochasticity of epithelial-immune interactions. In particular, the models show that the recruitment process of immune cells plays a pivotal role in driving successful immunosurveillance. However, this dependence involves several factors that remain undetectable from patient features, posing a challenge to the conventional process of data stratification. Subsequently, we identified a subset of virtual patients characterized by minimal outcome uncertainty, where cancer immunosurveillance failed. Here, the concept of digital twin templates becomes potentially effective. These templates group virtual patients with similar features and predicted trajectories, allowing for a more standardized approach to exploring treatment options. Within these templates, we simulated a hypothetical personalized immunotherapy in a controlled computational environment. This eliminates the inherent biological variability present in real-world patients. Successful therapeutic outcomes were achieved in a considerable portion of this patient subset, although a proportion experienced no discernible effects. Overall, this work provides important insights into the challenges associated with building CPDTs, including significant uncertainties in immune responses, challenges with data stratification, and the unpredictable nature of personalized treatments.

## 2 Results

### 2.1 The model overview of immune surveillance in micrometastases

The general dynamic of the model results from a series of rules that characterize crucial processes of immune-cancer cell interactions. We assume that the tumor microenvironment (TME) in the epithelial tissue is compounded by cancer, immune, and parenchymal cells. Cancer cells proliferate uncontrollably due to their high reproductive capacity, causing mechanical stress in the region of colonies. This mechanical stress leads to the death of adjacent parenchymal cells in the region, resulting in tissue damage and stimulating the activation of the immune system. The damaged tissue has a high concentration of cellular debris that stimulates the infiltration of macrophages and dendritic cells into the region. Moreover, macrophages and dendritic cells are activated through phagocytosis and contact, respectively. Macrophages carry out the phagocytosis process by ingesting dead cell waste and releasing tumor necrosis factor (TNF), a cytokine used for cell signaling, which leads to the recruitment of more immune cells to the region. This describes the transition process from the non-activated macrophage phase (M0) to the pro-inflammatory phase (M1). Dendritic cells (DCs) are activated when in contact with dying cells (cancer or parenchymal cells) and process their antigen material. For simplicity, we consider that all cancer cells present the same antigen. Activated dendritic cells migrate to the lymph node and present the antigen on their surface to T cells, promoting the activation and proliferation of helper and cytotoxic T cells. Consequently, the lymph node sends CD8+ (cytotoxic) and CD4+ (helper) T cells to the tumor microenvironment. CD8+ T cells perform two functions in the tumor microenvironment: (i) they kill cancer cells when they come into contact with them and (ii) inhibit the production of TNF from polarized macrophages, representing the conversion of classically activated macrophages (M1) to the anti-inflammatory state (M2) [34]. CD4+ T cells further enable the hyperactivation of macrophages, promoting their ability to engulf dead cells and live cancer cells [35] (Figure 1, more details in Methods Sections 4.1 to 4.7).

**Fig. 1:**
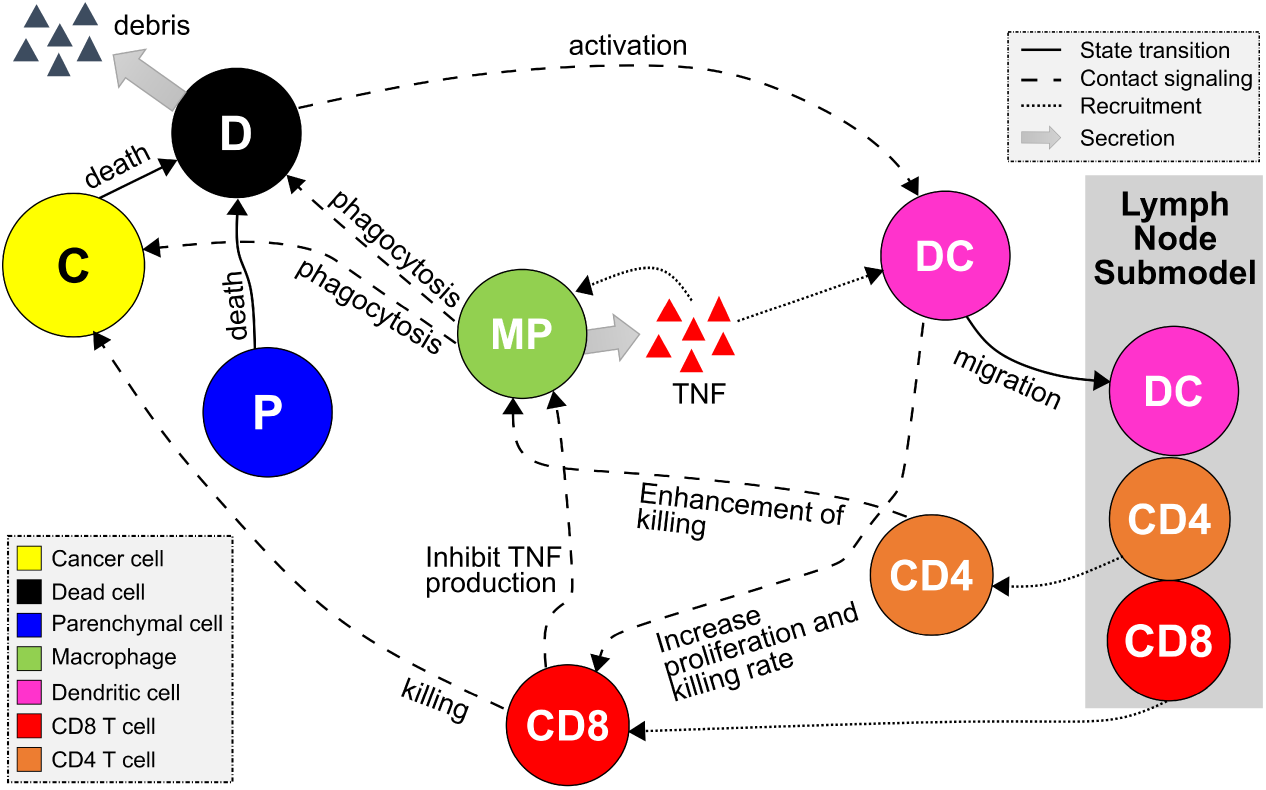
Schematic diagram of the multiscale model of immune surveillance in micrometastases. The contact of dendritic cells with the dying cells and the phagocytosis process by macrophages activate the immune response. Once the immune system is activated, the antigen presentation by DCs in the lymph node results in T Cell recruitment, which may extinguish cancer cells. Macrophages potentialize the immune response by releasing pro-inflammatory cytokines that increase the recruitment of immune cells.

### 2.2 Plausible patient outcomes

To investigate immunosurveillance in response to micrometastases growth, we explore a wide range of overall dynamics by adjusting various model parameters. Note that each parameter set can be seen as a distinct virtual patient, illustrating the spectrum of potential outcomes in virtual patient scenarios. In specific parameter configurations (additional information available in Section 1 of the supplementary material), we observed varying degrees of immune response during the progression of cancer.

We begin with a common initial state for all virtual patients. In this state, our model represents a region of epithelial tissue primarily composed of parenchymal cells, inactive immune cells, and a small number of metastatized cancer cells. These cells are randomly distributed in the microenvironment, as shown in Figure 2a. We then analyze the patient’s trajectories over a ten-day period, examining the variability in the responses of the immune system to micrometastatic development. Figure 2b demonstrates the simulation outcomes for specific virtual patients, highlighting three distinct intensities of immune response over time.

**Fig. 2:**
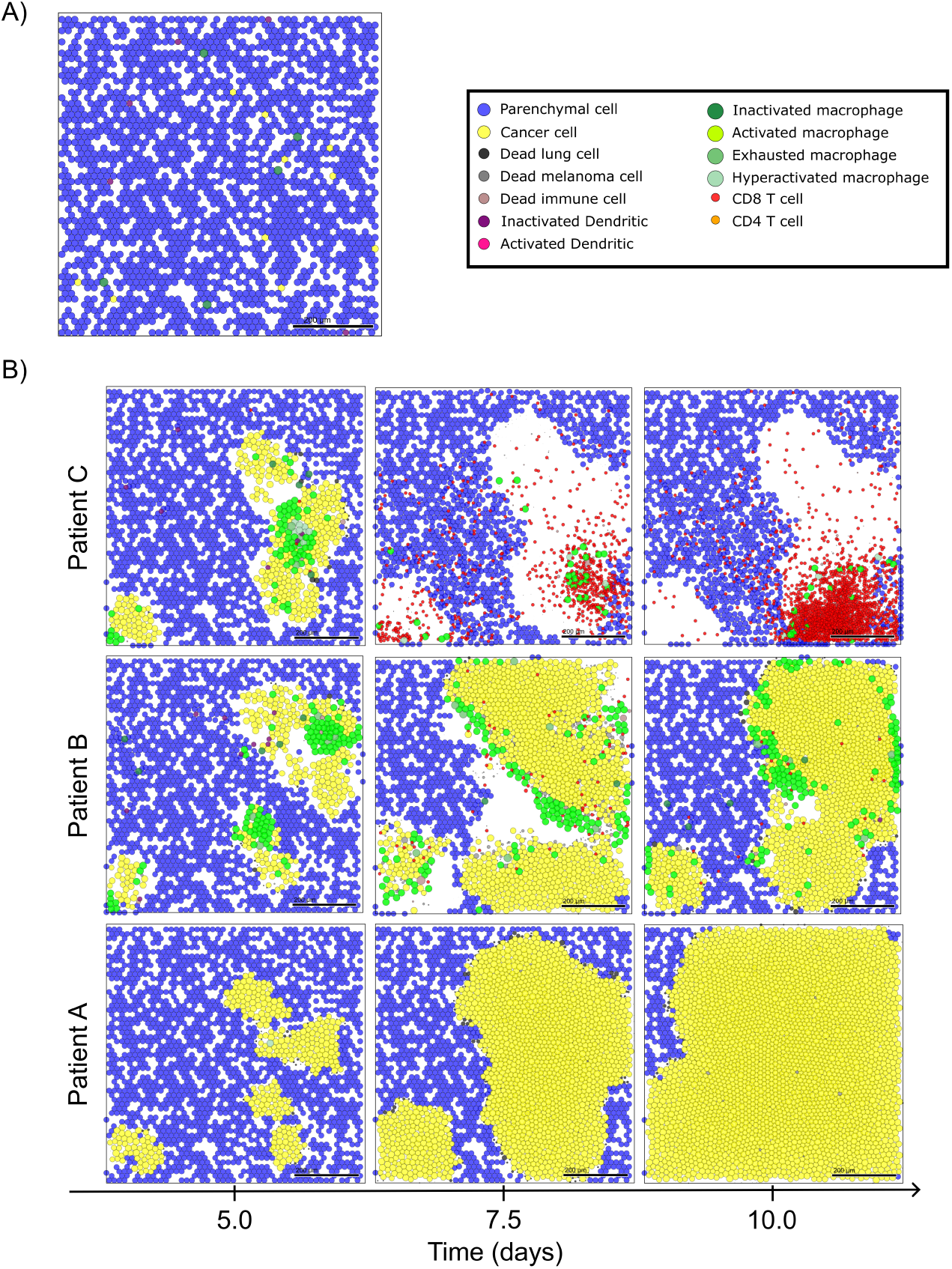
Multiscale model outcomes. A) Initial condition of multiscale model and legend of cells. B) Simulation snapshots under varying levels of immune response. From left to right, the days 5, 7.5, and 10 of the dynamics.

For the patient exhibiting a low immune response (Patient A in Fig. 2b), we observe a typical scenario of tumor escape, marked by the absence of immune cells and the prevalence of cancer cells in the microenvironment. The cancer progression exerts mechanical pressure on the surrounding tissue, resulting in the death of parenchymal cells and the creation of space for cancer cell proliferation. The stiffness of the tissue acts as a constraint on cancer progression, preventing excessive cellular agglomeration. However, the absence of macrophages precludes TNF production in the microenvironment, thus preventing the immune system activation and the production of adequate T cells required to combat the tumor.

In contrast, in the case of the patient displaying a moderate immune response (Patient B in Fig. 2b), we observe a scenario of partial elimination. This is characterized by the presence of active macrophages inducing TNF production and recruiting additional immune cells, leading to substantial immune control. However, the effective immune response occurs late, coinciding with significant damage to the tissue. This delayed response may be attributed to suboptimal DC activation or T cell production in the lymph node.

For the patient with a high immune response (Patient C in Fig. 2b), we observe a scenario of complete elimination, where the immune system effectively controls tumor growth. Early on, CD8+ T cells dominate the microenvironment, offering significant resistance against cancer cells. Furthermore, due to early immune activation, the tumor is eradicated before the ten-day simulation period concludes.

### 2.3 Virtual patient classification exposes limitations in conventional stratification

Utilizing virtual patient simulations generated from our parameter exploration (see Methods Section 4.8), we classified all 100,000 trajectories according to the immune response (see Figure 3a). To categorize the outcomes, we defined thresholds, *T_NC_* and *T_SC_*, for the population of live cancer cells in micrometastases at the end of the simulation (*P_M_*). These thresholds were established based on estimate carrying capacity of the computational domain, categorizing simulations as: tumor escape (no control, *P_M_ > T_NC_*), tumor elimination (significant control, *P_M_ < T_SC_*), and partial elimination (marginal control, *T_SC_ ≤ P_M_ ≤ T_NC_*). This classification resulted in 41.8% of simulations being categorized as significant control (SC), 45.1% as no control (NC), and 13% as marginal control (MC). Additional details and analysis of this classification can be found in Methods Section 4.9 and Section 4 of the supplementary material.

**Fig. 3:**
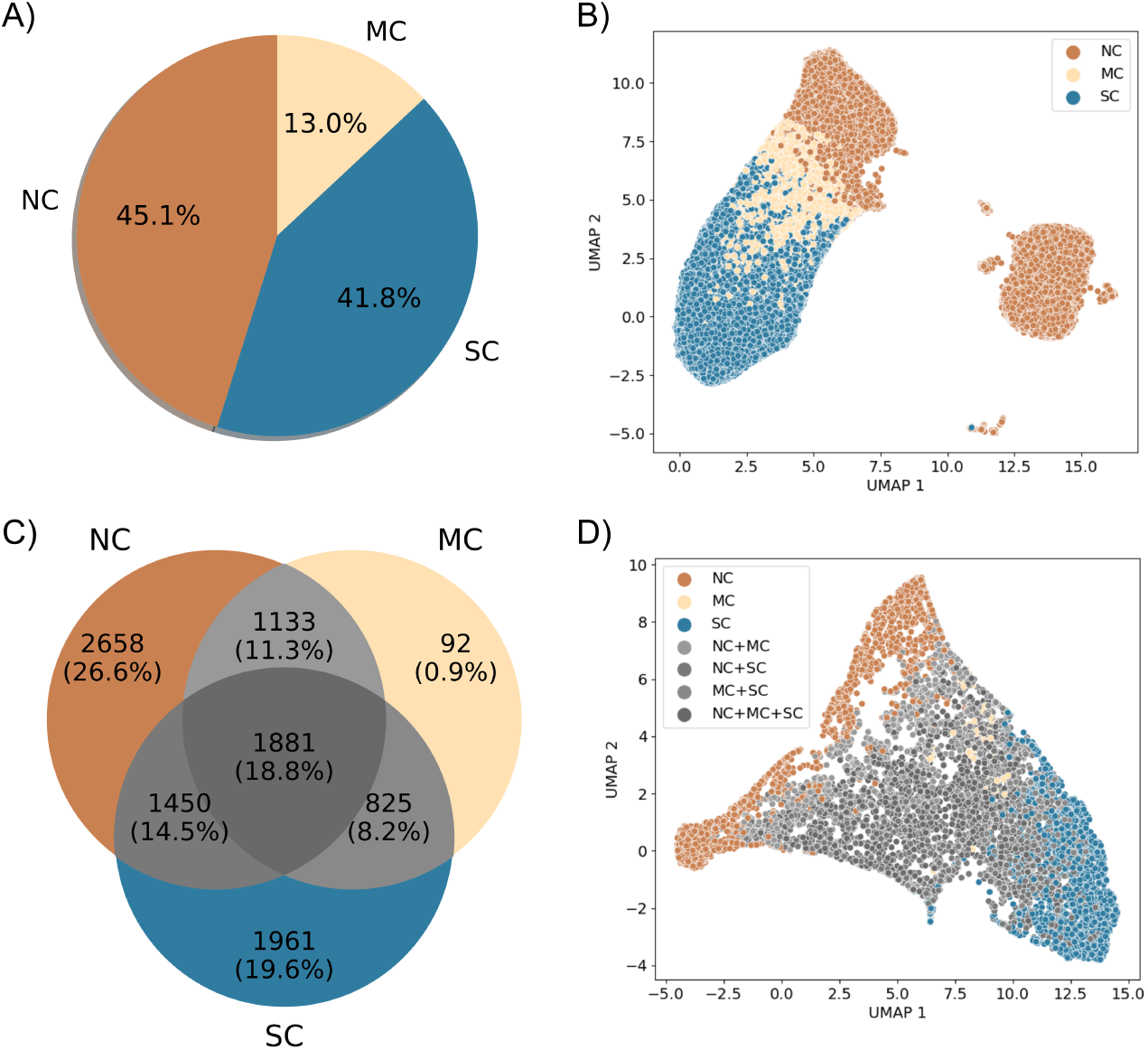
Virtual patient classification. NC: No control; MC: Marginal control; SC: Significant control. A) Classification of all 100,000 simulations according to the thresholds. B) Dimensionality reduction (UMAP) of the spatial/temporal trajectories of all simulations. C) Venn diagram of classification of the 10,000 virtual patients based on the totality of replicates. D) Dimensionality reduction (UMAP) of the averages of virtual patients trajectories.

After reducing the dimensionality of all 100,000 trajectories – each composed of 31 snapshots of RGB images (with dimensions of 5 *×* 5 *×* 3), resulting in a space of 2,325 dimensions – we observed two well-defined clusters. One of these clusters contains the three classes (NC, MC, SC) separated into distinct regions, while the other predominantly consists of trajectories classified as no control (NC) (Figure 3b). We further investigated the differences between these two clusters of NC trajectories and found evidence suggesting that they differ in terms of immune system activation. The cluster containing all classes has an activated immune system, whereas the other one is deactivated or in the process of deactivation (more details in Section 3 of the supplementary material).

Conflicts arose in the classification of replicates for individual virtual patients. This is because the stochastic nature of immune-tumor interactions introduces substantial uncertainty in the outcomes. Notably, out of the 10,000 virtual patients, only 47.1% exhibited consistent classification for all replicates within the same category (Figure 3c). For the remaining patients (52.9%), a unique outcome label was not assigned uniformly across all replicates.

When classifying patients based on the entirety of replicates and visualizing the mean trajectories in a reduced space (each mean trajectory is calculated from 10 replicates, giving a total of 10,000 mean trajectories), we observed that mainly NC and SC patient labels preserve distinct regions in the reduced space. Patients with multiple labels (NC+MC, NC+SC, MC+SC, and NC+MC+SC) and those with the MC label are intermingled in the reduced space (see Figure 3d). This is because of the lower variances in the immune responses of patients with extreme labels: NC and SC. On the other hand, for some patients, their outcomes have significant variance. For instance, patient 6 (Figures 4a and 4b) displays sources of instability in the immune response and concurrently exhibits the three labels SC, MC, and NC. In contrast, patient 5 (Figures 4c and 4d) has a unique significant control label across all replicates. This observed diversity in these patient’s outcomes can be attributed to substantial variations in immune cell population across different replicates, particularly noticeable with the population of CD8+ T cells, dendritic cells (DCs) and macrophages (Figures 4b and 4d).

**Fig. 4:**
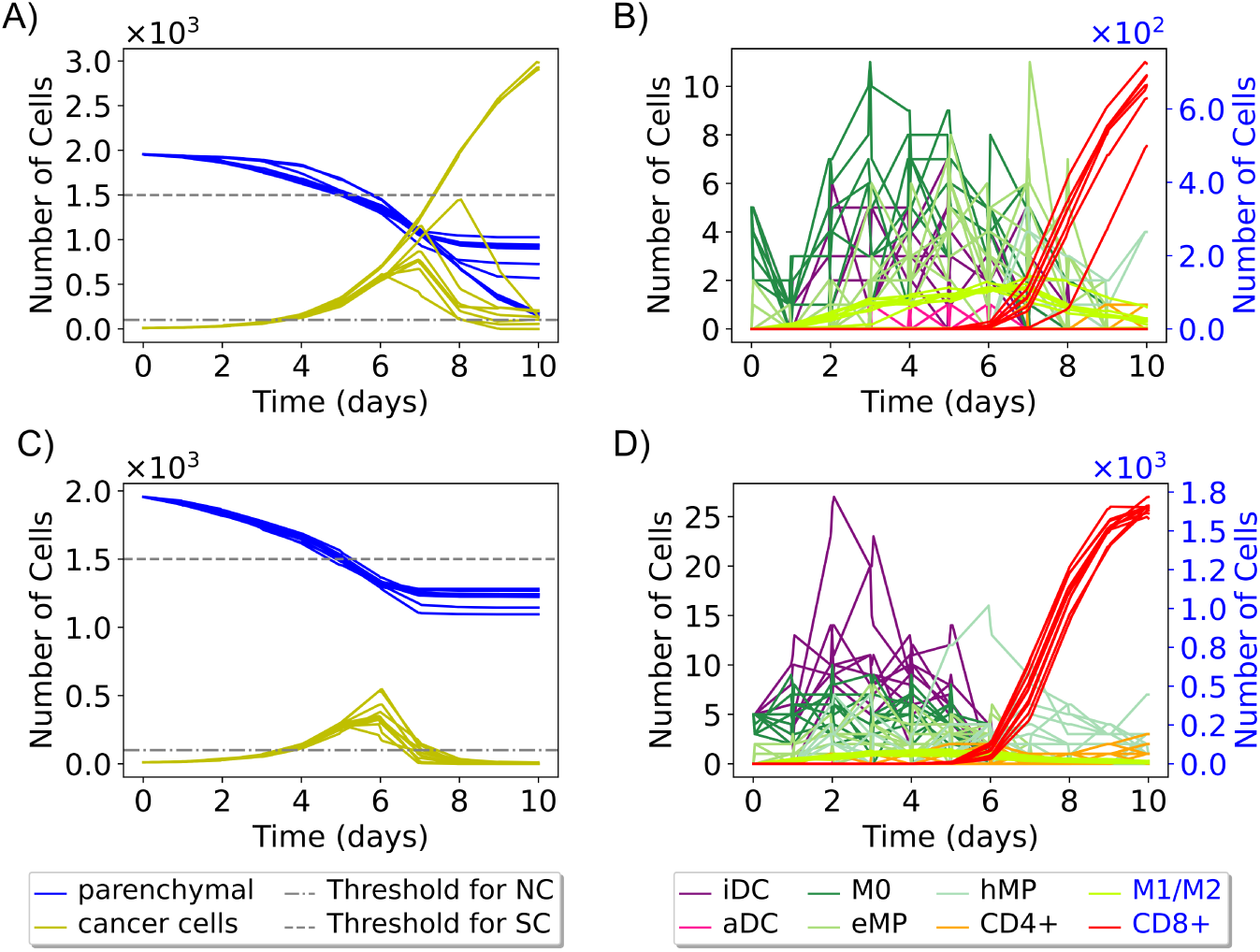
Cell Population Trajectories for Virtual Patients. Ten trajectories show the change in cancer, parenchymal, and immune cell populations over time for two virtual patients. Patient 6 (panels A, B) exhibits various outcomes (NC, MC, SC) across replicates. Patient 5 (panels C, D) has a unique “significant control” (SC) outcome. (iDC: inactive dendritic cells, aDC: active dendritic cells, eMP: exhausted macrophages, and hMP: hyperactivated macrophages)

In the proposed model, variations in immune responses are primarily driven by the recruitment effect. Early recruitment leads to significant immune responses, whereas later recruitment is often associated with weaker or non-existent responses. For instance, in Figure 4b, the curves for macrophages M0 and inactive dendritic cells (iDCs) show a decay between day 0 and day 2, followed by an increase, indicating that these cells are recruited later, contributing to a more substantial immune response. Conversely, in Figure 4d, where there is no observable decay, it suggests that M0 macrophages and iDCs are recruited early, resulting in a more pronounced immune response. In addition to the recruitment effect, increasing the populations of dendritic cells and macrophages is crucial for activating the immune system. This activation leads to the recruitment of CD8+ T cells. The activation of dendritic cells triggers their migration to the lymph node, where they initiate the activation and differentiation of T cells into CD8+ and CD4+ T cells. CD8+ T cells then eliminate cancer cells, as reflected in the varying cancer cell population trajectories observed in Figures 4a and 4c.

Additional information on these patients’ trajectories and others is provided in Table 7 of the supplementary material. The timing of immune cell recruitment – whether early or late in tumor progression – has a significant impact, consistent with established biological findings [36, 37].

Given the variability in patient outcomes, the conventional approach to patient stratification proves inadequate for all virtual patient datasets. Consequently, there arises a requirement for additional information on patient trajectories to mitigate uncertainty in immune responses, with a specific focus on elucidating the complexities of the immune cell recruitment process. Our future investigative endeavors aim to incorporate such detailed mechanisms and integrate multiple data sources into our model for a comprehensive understanding.

### 2.4 Sensitivity analysis reveals key patient characteristics

In the parameter space of the virtual patients, we identify the locality, spread, and skewness of the ten parameters according to the labeled patient trajectories (see Figure 5a), and perform a statistical hypothesis test to examine the relationship between the classes: No Control (NC) and Significant Control (SC). Considering as null hypothesis that the distribution of each parameter is the same for both classes, we performed the Mann-Whitney U test, according to a confidence level of 95%, we obtain that the parameters *r_leave_* (rate of activated DCs leaving the tissue and migrating into the lymph node) and *δ_C_*(natural decay of T cells on lymph node) are not statistically different in either class (p-value *≥* 0.05), while the others parameters are statistically different (p-value *<* 0.05). However, the parameters *r_recruit_*[*D*] (recruitment rate of dendritic cells) and *r_recruit_*[*M*] (recruitment rate of macrophages) have greater statistical significance in our parametric space. This suggests that the process of recruitment of macrophages and dendritic cells is a crucial component of our model. Their strong associations with patient labels and the outcomes further support our findings regarding the specific virtual patient examined in the previous Section 2.3.

**Fig. 5:**
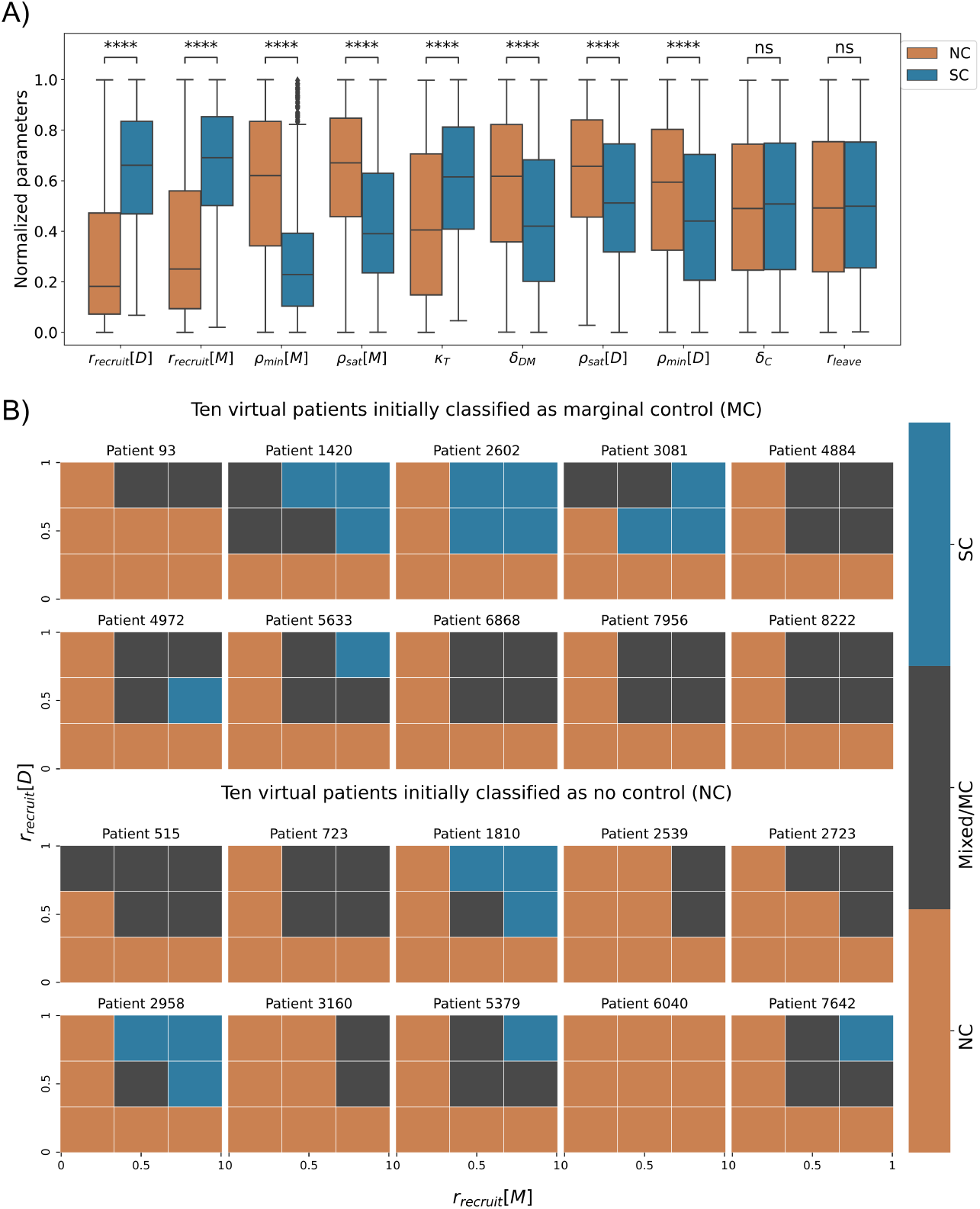
A) The quartiles of normalized parameters according to each label. The box bounds define interquartile ranges (IQR). The line that divides the IQR represents the median, and the whiskers’ length is 1.5 times the interquartile range above the upper quartile and below the lower quartile (parameters in ascending order of p-value, ****: p-value *<* 10*^−^*^4^ and ns: p-value *≥* 0.05). B) Analysis of 20 patients on normalized parametric space *r_recruit_*[*M*] *× r_recruit_*[*D*]. The first two rows are 10 patients labeled as MC, and the bottom 2 rows are 10 patients labeled as NC. The mesh of parametric space is regular, and 3 *×* 3 for each patient, the color represents the label of mesh partition.

### 2.5 A hypothetical personalized immunotherapy

According to the multiscale model developed, the recruitment of macrophages and dendritic cells emerges as a pivotal factor in orchestrating the antitumor response. Changes in dendritic cell recruitment rates are associated with various therapeutic modalities, including immunotherapy, radiotherapy, and chemotherapy, resulting in distinctive modulation of the immune response [38]. The classification of macrophages is intricate, traditionally relying on the M1 and M2 phenotypes. The M1/M2 ratio serves as a consequential prognostic indicator, where a diminished value denotes a pro-tumor microenvironment [39]. Notably, within our model, macrophage recruitment positively influences the patient’s prognosis. However, in a broader context, macrophage recruitment is commonly associated with tumor survival, and inhibiting this process is considered a potential therapeutic avenue [39]. The observed incongruence with our model may be explained by the absence of supplementary anti-inflammatory dynamics activated by tumor-associated macrophages (TAM).

We performed a hypothetical immunotherapy in twenty virtual patients, where we applied changes in the parameter space *r_recruit_*[*M*] *× r_recruit_*[*D*] to test the impact on clinical outcomes. Ten of these patients were randomly sampled from the 92 patients labeled as MC and ten from the 2658 patients labeled as NC (Figure 3b). For each sampled patient, we performed simulations using the possible combinations of minimum, central, and maximum values of each of the two parameters, defining the 3×3 mesh. In each partition, we execute ten replicates and label the outcomes according to our classifier (details in Methods Section 4.9), resulting in a total of 90 simulations for each patient (Figure 5b, generated from 1800 new simulations).

In Figure 5b, the results show overall that the higher the recruitment rates, the greater the chance of the patient to have a better response from the immune system, in some cases being able to generate a significant control of tumor growth, for example, patients 1420, 2602, 3081, 4972, 5633, 1810, 2958, 5379, and 7642. The patients 3081 and 4972 outcomes show that the relationship between recruitment rate of DCs and SC label is not entirely linear. On the other hand, for some patients, even varying recruitment rates do not change the outcome, for example, patient 6040. This fact highlights the significance of integrating patient-specific treatment with mechanistic models for treatment efficacy prediction. Mechanistic models empower us to identify patient-specific characteristics and assess treatment alternatives.

## 3 Discussion

In this study, we build a multiscale model to represent early micrometastases growing in general epithelial tissue. We modeled the abnormal cancer cell proliferation generating a cascade of the effects that trigger the immune system response. The growth of the micrometastases creates mechanical stress in healthy tissue, inducing pro-inflammatory signals. The signals attract and recruit more immune cells to the metastatic region creating cancer immuno-surveillance. The model is able to generate different immune control responses including escape, and partial and complete elimination. Next, we used high performance computing for large scale exploration of ten model parameters, focusing on those that modulate the immune response. We used Latin Hypercube Sampling to select 10,000 virtual patients (each patient is characterized by a unique set of parameter values), and performed 10 replicates for each patient, generating a total of 100,000 simulations. Furthermore, based on thresholds for the cancer cell population after 10 days of the simulation, we classified the outcomes into three distinct behaviors of tumor control: significant control (SC), marginal control (MC), and no control (NC). Our model successfully represents two of the three Es of cancer immunoediting [40], namely elimination and escape, through the simulation of these scenarios. To incorporate the concept of equilibrium, we propose the integration of additional phenomena into the model, such as the healing process of damaged tissue and anti-inflammatory responses.

The dimensionality reduction of trajectories in spatial-temporal data supports the distinction of three separate behaviors. Furthermore, the reduced space reveals a distinct set of trajectories suggesting a unique subtype of no-control behavior (NC). We have demonstrated that the differences between these two NC clusters are associated with immune activation. The reduced space of trajectory averages further underscores the significant uncertainty surrounding certain patient outcomes and effectively separates patients into either NC or SC labels. Moreover, this model reduction process may not represent the optimal approach for spatial-temporal trajectories generated by the model. Additional alternatives should be considered for evaluation in the near future [41–44].

After analyzing the parameter space of virtual patients, we have identified that the recruitment process of macrophages and dendritic cells has the most significant impact on tumor control by the immune system. Further exploration has suggested that altering the values of key parameters in certain patients can lead to changes in outcomes. While this variation in patient labeling is purely computational, in a scenario where the model is validated, this study can be interpreted and linked to potential therapeutic strategies for specific patients. However, we emphasize that our patient classifier is preliminary, and the development of a trajectory classifier that considers space/time variations and the stochasticity of the model is the subject of future investigations [12].

The model developed and implemented in this work focused on early metastatic events and thus only a subset of relevant immune interactions were considered. We are currently working on extending the model to include additional interactions such as the variability of tumor specific antigens, which can affect the capacity of the immune system to detect and kill metastatic cancer cells; this will be used to simulate the efficacy of antigen-specific treatments. Moreover, we plan to extend the model by adding more interactions and cell types, including cells and cytokines that drive an anti-inflammatory response. We will also extend the model to include cells of the innate immune response, such as NK cells, and also cells that are known to greatly affect the microenvironment and the immune response such as cancer-associated fibroblasts (CAFs).

The immediate future path for this work involves using the model to identify digital twin templates, which are groups of virtual patients with similar characteristics and predicted trajectories, allowing for a more standardized approach to exploring treatment options [11, 12]. However, during this work, we have encountered challenges along this path. Based upon our work, we find that the concept of creating patient templates would fail for a considerable number of patients: simulations show that even with perfectly known parameters values and initial conditions, the inherent stochasticity of immune-tumor interactions can drive radically divergent trajectories from similar (or identical) initial conditions. For such patients, additional post-calibration measurements would be required to drive the twin towards the patient’s actual trajectory. On this basis, we recognize the necessity of obtaining more spatial and temporal information from untreated patients. This type of data would allow for a more comprehensive characterization of the initial tumor and immune state, with personalized model refinement whenever new patient measurements come available (e.g., during post-treatment care). This combination of refined initial characterization with integration of new patient measurements could lead to the development of more accurate digital twins. However, obtaining such detailed information is often a challenge in typical clinical settings due to the urgency of initiating treatment for patients. To overcome these challenges, our plan is to integrate real-time molecular data and calibrate time-course data acquired from wet labs into a comprehensive agent-based model. This model can then be translated into a more general model to predict patient outcomes. With this novel approach, we can address two crucial aspects: A) Predicting the long-term trajectory of a patient by integrating multiple data sources, instead of relying solely on initial time data, which can lead to excessively variable outcomes; B) Continuously assimilating subsequent patient data, which is essential for accurately discerning prognoses of patients. Consequently, maintaining an ongoing alignment with new patient measurements will empower the Cancer Patient Digital Twin (CPDT) to anticipate the future disease state and forecast its response to treatment [15, 45]. This transformative strategy equips clinicians to select the most suitable treatment for each patient, revolutionizing cancer care by enhancing patient outcomes while minimizing adverse effects.

## 4 Methods

### 4.1 Multiscale model construction

We developed the multiscale model using PhysiCell [46], an open-source framework that allows the development of multicellular models at various spatial/temporal scales. In this tool, the cells are represented by off-lattice agents with independent behaviours, including cell cycle progression, death processes, volume changes, and motility. Some of these dynamics may be defined by substrate availability in the environment, which is represented on PhysiCell using an open-source biological diffusion solver, BioFVM [47]. PhysiCell has been used in a wide variety of multicellular problems, such as virus therapy, cancer immunology, tissue mechanics, and drug screening, among others [16, 23, 48–50].

We have previously formulated a model that elucidates the immune response to the dissemination of SARS-CoV-2 in the lungs [23]. This simulator, designed to emulate SARS-CoV-2 interactions within lung tissue, incorporates a collection of mechanisms that are based on domain expertise in immunology, virology, and tissue biology. In the current study, we have adapted this model to investigate the emergence of micrometastases in general epithelial tissue. The cellular components of the tumor microenvironment are formed by parenchymal, immune, and cancer.

### 4.2 Micrometastasis and epithelial tissue interaction

We modeled parenchymal cells in homeostasis in the absence of tumor cells. As the tumor grows, cancer cells create intense mechanical pressure on the tissue, causing extensive damage to the region, eventually leading to the death of healthy cells [51]. Similar to [49], we model elasto-plastic mechanics of parenchymal cells following adhesion-repulsion mechanics [46, 52], and interactions associated with an adjacent extracellular matrix [53]. Specifically, cells are subjected to two forces i) elastic force that resists their displacement and ii) plastic reorganization where the cell adheres to another point of the extracellular matrix, generating a rearrangement process. Specifically, given that cell *i* with position *x_i_* and radius *R_i_* is attached to the extracellular matrix at 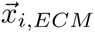 experiencing the two forces described above and considering 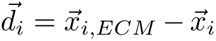 and 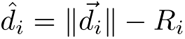 to be very small, we have

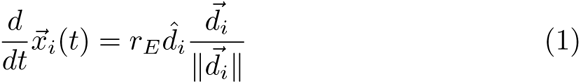

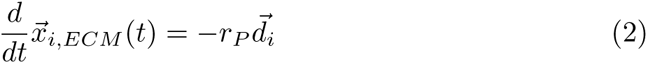

where *r_E_*is a coefficient for the magnitude of an elastic restorative force and *r_P_* is a relaxation rate. If 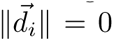, the cell is in equilibrium, implying 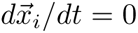 and 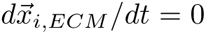. This mechanical stress generates two effects on the dynamics between parenchymal and cancer cells. The parenchymal cell undergoes programmed cell death if it has a deformation induced by a cancer cell greater than a maximum deformation value (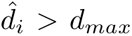), providing cancer cells with additional space to proliferate. On the other hand, cancer cells suffer a decreased proliferation capacity due to the mechanical pressure sustained by the tumor microenvironment [49]. Based on a cell cycle model (flow cytometry separated) from PhysiCell [46], we define the probability of a cell transitioning from phase G0/G1 to phase S in a time interval Δ*t* depending on the local compressive pressure, given by:

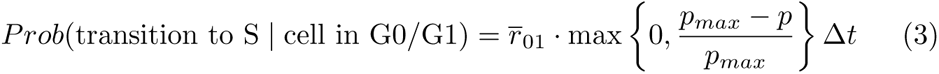

where *p* is the simple pressure (a dimensionless analog of mechanical pressure [46]), 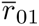 is the rate in the ideal mechanical situation for a cell to start the proliferation process, and *p_max_* is the simple pressure threshold at which cancer cells stop proliferating.

### 4.3 Motility and infiltration of immune cells

The immune cells migrate into the tumor microenvironment via chemotaxis, guided by the tumor necrosis factor and debris gradients secreted by macrophages and dying cells, respectively. Cell motility is defined by their speed, bias direction, and persistence time in the velocity direction (*T_per_*) as described in [46]. Mathematically, we calculate the velocity by

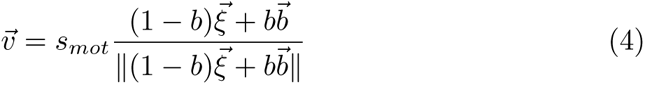

where *s_mot_* is the speed of chemotaxis, *b* denotes the migration bias (with a range of 0 to 1), 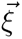 is a random unit vector direction, and 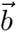 is the migration bias direction. Velocity undergoes stochastic changes in accordance with a Poisson probability distribution with a rate 1*/T_per_*, as discussed in [46]. In this study, we consider 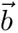 as a combination of tumor necrosis factor and debris gradients, defined by the equation

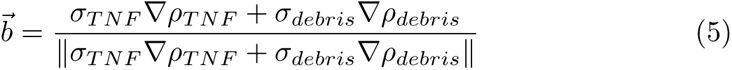

where *σ_T NF_* and *σ_debris_* are the sensitivity of chemotaxis along with the TNF or debris gradient, respectively. We assume that macrophages and activated dendritic cells (DCs) exhibit TNF and debris sensitive chemotaxis (*σ_T NF_ >* 0 and *σ_debris_ >* 0), while non-activated DCs and macrophages rely on the debris gradient for chemotaxis (*σ_T NF_* = 0 and *σ_debris_* = 1). CD8+ and CD4+ T cells migrate exclusively based on the contribution of the TNF gradient (*σ_T NF_* = 1 and *σ_debris_* = 0). Activation of DCs and macrophages reduces motility speed, while cells attached to other cells remain immobile.

Eventually, immune cells infiltrate the tumor microenvironment through the vasculature. In this model, we randomly sampled voxels within the computational domain and designated them as vascularized voxels. These vascularized voxels are assumed to represent approximately 8.8% of the total tissue volume [23, 54]. Immune cells recruited to the tumor microenvironment infiltrate the domain through these vascularized voxels. Macrophages and dendritic cells are recruited to the microenvironment due to high concentration of TNF, in a time interval Δ*t*. Cells are recruited to the system according to the following equation:

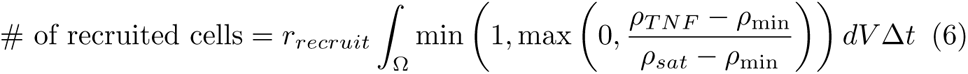

where *r_recruit_* is the recruitment rate, *ρ*_min_ and *ρ_sat_* are the minimum and maximum signal of TNF to recruit the cells, *ρ_T NF_* is the TNF concentration in each partition of the domain Ω, and *dV* is the volume element of Ω. CD8+ and CD4+ T cells are recruited to the tissue based on the response of the lymph node after dendritic cell migration into the lymph node [23].

### 4.4 Phagocytosis and macrophage phases

Macrophages play a key role in the dynamics of cancer and the immune system, which we capture in this model [55]. They carry out the cleaning process of the tissue by the process of phagocytosis (engulfment) of dead cells, which triggers the production of pro-inflammatory signals (in our model, TNF release) [56]. Eventually, inactivated macrophages (M0), attracted by debris released by dying cells, phagocytize the dying cell following a conditional probabilistic event within a time interval Δ*t*:

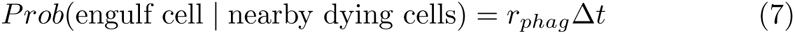

where *r_phag_*represents the phagocytosis rate of the macrophages. In this process, macrophages become activated (M1, classically activated, proinflammatory) and absorb the entire volume of the dying cell (Figure 6a). M1 macrophages spend time ingesting the genetic material and are unable to phagocytose during this phase. Moreover, the M1 macrophages reach exhaustion by ingesting a large amount of cell debris, leading to the loss of their ability to phagocytose cells and increasing the likelihood of their own death [57] (Figure 6b). Activated macrophages in contact with CD4+ T cells become hyperactivated, enhancing their potential to phagocytose cells, including the possibility of engulfing live cancer cells [34] (Figure 6c). The CD8+ T cells, when in contact with M1 macrophages, inhibit the production of TNF by the macrophages [34] (Figure 6d).

**Fig. 6:**
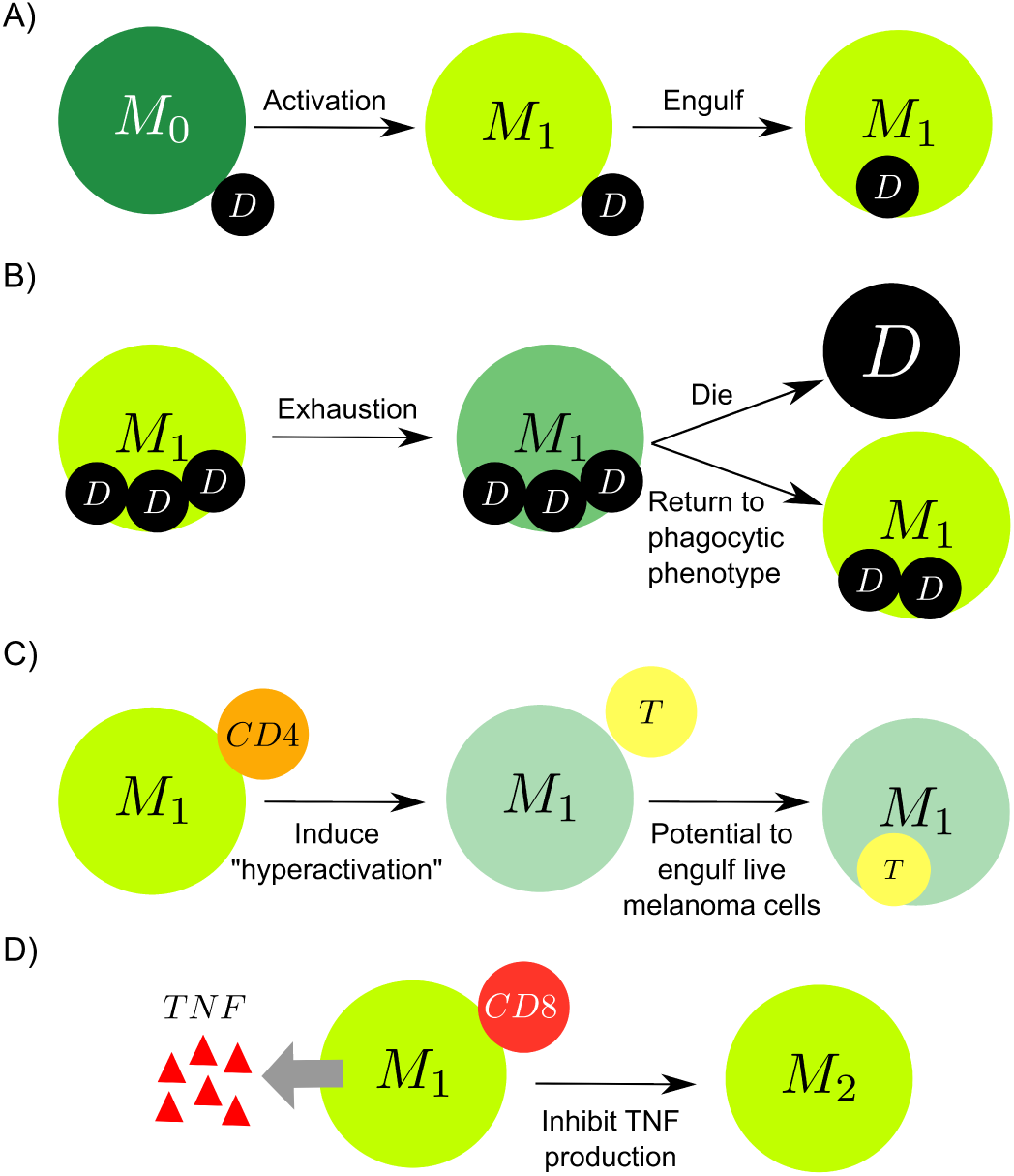
Phagocytosis and macrophage phases. A) Macrophages in contact with dead cells are activated (M1 phase) and engulf the dead cell. B) M1 macrophages reach exhaustion by engulfing above a certain threshold of debris and stopping phagocytosing, increasing the chance of dying and decreasing the possibility of returning to the phagocytic phenotype. C) The hyperactivation of macrophages induced by CD4+ T cells increases the potential to phagocytose live cancer cells. D) Activated macrophage in contact with a CD8+ T cell activates an anti-inflammatory response (M2) and stops secreting TNF.

### 4.5 Dendritic cell behavior

Initially, all dendritic cells (DCs) are non-activated in the tissue. The DCs attach cancer or a dying parenchymal cell and become activated, representing the process of antigen presentation and inflammatory mediators produced by dead parenchymal cells from mechanical stress. The attachment process is modeled in a time interval *δt* according to the following probabilistic event:

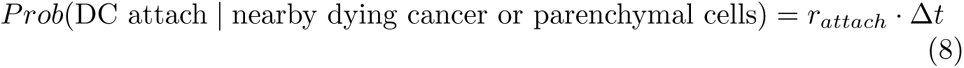

where the *r_attach_* is the attachment rate of DCs. Activated DCs migrate to the lymph node according to a probabilistic event with a rate *r_leave_*:

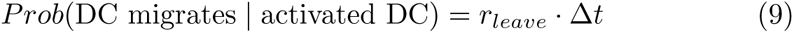

in any time interval [*t, t* + Δ*t*]. The migration of DCs to the lymph node spend a time *τ_DC_*. In the lymph node, these cells trigger the production, proliferation, and activation of specific T cells (details in Methods Section 4.6). In the tumoral microenvironment, activated DCs enhance the CD8+ T cell proliferation capability and attachment rate (details in Methods Section 4.7).

### 4.6 Lymph node submodel

We defined the lymph node model from the time course of cell populations. The number of dendritic cells that have migrated from the tumor microenvironment to the lymph node (*D_M_*) upregulates the proliferation and activation of two types of helper T cells (*T_H_*_1_ and *T_H_*_2_) and cytotoxic T cells (*T_C_*). Consequently, these cytotoxic T cells activate the production of CD8+ cytotoxic (*T_Ct_*) and CD4+ helper (*T_ht_*) T cells, which will be trafficked to the tumor microenvironment [23, 58–60]. Altogether, the variation of the cell populations over time is given by:

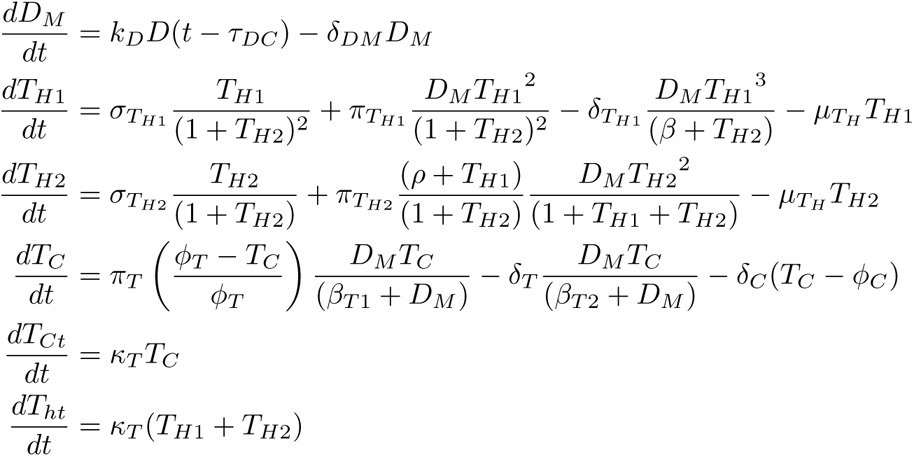

where *k_D_* and *τ_DC_* are the antigen presentation rate by dendritic cells and the time duration by the DCs to migrate into the LN, *δ_DM_* is the natural decay of *D_M_*, *σ_TH_*_1_ and *σ_TH_*_2_ denote the proliferation rates for type 1 and type 2 helper T cells, *π_TH_*_1_ is the activation rate of *T_H_*_1_ induced by *D_M_*, *δ_TH_*_1_ and *β* are rates and half maximum deactivation mediated by *D_M_* for type 1 helper T cells, *µ_TH_* is the natural death rate of both cell types *T_H_*_1_ and *T_H_*_2_, *π_TH_*_2_ represents the activation rate of type 2 helper T cells mediated by *D_M_*, *ρ* is a weight conversion between *T_H_*_1_ and *T_H_*_2_ activation, *π_T_* and *δ_T_* are activation and deactivation of *T_C_* mediated by *D_M_*, *φ_T_* is scaling factor between lymph node and tumor microenvironment, *β_T_* _1_ and *β_T_* _2_ are half maximum of activation and deactivation of T cells mediated by *D_M_*, *δ_C_* and *φ_C_* are the natural decay and population threshold of T cells, *κ_T_* is the recruitment rate of cytotoxic and helper T cells. The CD8+ and CD4+ T cells infiltrate the tumor microenvironment through the vasculature after a time *τ_T_*, according to the cells amount *T_Ct_*(*t − τ_T C_*) and *T_ht_*(*t − τ_T C_*), respectively [23, 61, 62]

### 4.7 Cytotoxic CD8+ T cells killing cancer cells

Given a cancer cell in the neighborhood of CD8+ T cells, the probability of CD8+ T cells attaching to this cell in a time interval Δ*t* is given by:

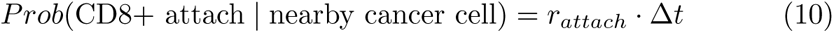

where *r_attach_* is the rate of attachment [16, 23, 63]. Once the cells are attached, they can eventually detach, according to a probabilistic event that is a function of the duration of attachment *T_attach_* as following:

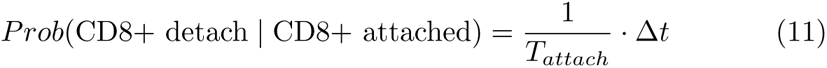

in any time interval [*t, t* + Δ*t*]. Otherwise, cells remain bound until the time *T_attach_*, when the cancer cell dies. Eventually, multiples CD8+ T cell can attach to the same cancer cell, increasing the likelihood of killing.

### 4.8 Generating virtual patients

Using HPC resources, we conducted a parameter exploration to generate virtual patients employing the multiscale model discussed earlier. Specifically, we identified ten parameters associated with the immune response, out of which six were linked to the recruitment of dendritic cells and macrophages to the microenvironment, three were associated with the production of T cells in the lymph node, and one was related to the migration of activated dendritic cells from the microenvironment to the lymph node. To sample the parameter space, we employed Latin hypercube sampling (LHS) and generated 10,000 parameter sets (see Section 1 in supplementary material). To account for the stochastic nature of the model, we created ten replicates for each parameter set, resulting in a total of 100,000 simulations. Due to the high dimensionality of the data, we compressed the data significantly to facilitate analysis (see section 2 in supplementary material).

The average wall time for each simulation, utilizing eight threads in shared memory programming with OpenMP, is approximately 11 minutes. If 100,000 simulations are processed in serial, it could take around 764 days. However, by utilizing HPC resources and executing 300 model simulations simultaneously (2400 threads), the entire dataset can be generated in 2.5 days.

### 4.9 Patient classifier

Based on the live cancer cell population for each simulation, we created a classifier to distinguish different immune system responses. We define two threshold values (*T_SC_* and *T_NC_*) to categorize three types of responses from the cancer cell population on day 10 (*P_M_*). The first type of response is significant control (SC),where most of the cancer cells have been eliminated by the immune system at the end of the simulation (day 10), indicated by *P_M_ < T_SC_* (elimination). The second type is no control (NC), where the immune system is not strong enough to eliminate the tumor, indicated by *P_M_ > T_NC_* (escape). The third type is marginal control (MC), an intermediate situation where there is partial elimination of the tumor, indicated by *T_SC_ ≤ P_M_ ≤ T_NC_* (partial elimination).

The thresholds were defined based on the estimated total cell population in the domain (approximately 2,000 cells), with *T_NC_* set at 75% of the total population (1,500 cells) and *T_SC_* set at 5% of the total population (100 cells). These initial thresholds were chosen to explore the classification landscape and require further validation. In Section 4 of the supplementary material, we compare this classifier strategy with a clustering method that utilizes a dedicated time series metric (dynamic time warping). This analysis, demonstrates that our classifier has low dependence on individual time points, particularly for the extreme cases of no control (SC) and significant control (SC), where our accuracy remains above 90%.

## Data availability

The dataset publicly is available in [64].

## Code availability

The multiscale model and postprocessed dataset publicly are available in https://github.com/heberlr/micromets_immuno/releases/tag/1.0.5.

## Acknowledgments

This work was supported in part by Cancer Moonshot funds from the National Cancer Institute, Leidos Biomedical Research Subcontract 21X126F. H.L.R., M.G., and P.M. were supported by Jayne Koskinas Ted Giovanis Foundation for Health and Policy, the National Science Foundation (Grants 1720625 and 2303695), and Luddy Faculty Fellowship. B.A. and I.S were supported by U.S. Department of Energy (DOE) (Grant DE-SC0021631). We thank Jeffrey Bryan (University of Missouri) and Tina Hernandez-Boussard (Stanford University) for their helpful discussions. The computational simulations of this research were performed on high-throughput computing clusters at Indiana University, which were supported in part by Lilly Endowment, Inc., through its support for the Indiana University Pervasive Technology Institute.

## Competing Interests

The authors declare no competing interests.

## Author contributions

H.L.R., B.A., I.S. and P.M. conceptualized and designed the study. H.L.R., B.A. and M.G. developed the computational model. H.L.R. built the database and performed computational and statistical analyses. I.S. and P.M. managed and supervised the project. All authors wrote and reviewed the manuscript.

## Notes

https://github.com/heberlr/micromets_immuno

### Competing Interest Statement

The authors have declared no competing interest.

### Summary of Updates

Introduction: Updated to include a more comprehensive description of CPDT, its limitations, and the connection with the modeling exercise conducted in this study. Results: Improved readability by refining figures, descriptions, and explanations. Discussion: Expanded to include detailed points on the limitations encountered in this study and the types of data that could help overcome these limitations. Supplemental Material: - Enhanced descriptions in Section 4 on data compression processes. - Added a new section, Section 4, with classification analysis. - Introduced a new Table S7, showcasing trajectories of four patients grouped by immune response.

