## Supplementary Material for "A multiscale model of immune surveillance in micrometastases: towards cancer patient digital twins"

#### 1 Parameters tables

The multiscale model that has been constructed contains hundreds of parameters. In Tables S1 to S3, we present the essential parameters defined for a hypothetical middle patient, which represents the average characteristics of all patients. The set of virtual patients was created based on ten parameters that are associated with the immune response. Out of these ten parameters, six are linked to the recruitment of dendritic cells and macrophages into the microenvironment ($r_{recruit}[MP]$, $\rho_{min}[MP]$, $\rho_{sat}[MP]$, $r_{recruit}[DC]$, $\rho_{min}[DC]$, and $\rho_{sat}[DC]$), three are associated with the production of T cells in the lymph node ($\delta_C$, $\kappa_T$, and $\delta_{DM}$), and one is related to the migration of activated dendritic cells from the microenvironment to the lymph node ($r_{leave}$). The parameter range values for generating the virtual patient dataset using Latin Hypercube Sampling (LHS) are shown in Table S4.

**Table S1**: Table of motility, substrate secretion, and recruitment parameters.

| Parameter | Description | Value | Unit | Reference |
|---|---|---|---|---|
| $\sigma_{TNF}$ | sensitivity of chemotaxis along with TNF gradient | 10 | – | Estimated |
| $\sigma_{debris}$ | sensitivity of chemotaxis along with debris gradient | 1 | – | Estimated |
| $s_{mot}$[DC, CD4+] | speed motility of DCs and CD4+ T cells | 2 | $\mu m/min$ | [1] |
| $s_{mot}$[M0, CD8+] | speed motility of M0, and CD8+ T cells | 4 | $\mu m/min$ | [1] |
| $s_{mot}$[M1,M2] | speed motility of M1 and M2 | 0.4 | $\mu m/min$ | [1] |
| $T_{per}$ | persistence time on velocity direction | 5 | $min$ | [1] |
| $b$[MP, DC, CD4+] | migration bias of DCs, MPs, and CD4+ T cells | 0.7 | – | [1] |
| $b$[CD8+] | migration bias of CD8+ T cells | 0.3 | – | [1] |
| $U_{TNF}$[DC,M0,CD8+] | TNF uptake rate of DCs, M0, and CD8+ T cells | 0.01 | $min^{-1}$ | Estimated |
| $U_{TNF}$[M1,M2] | TNF uptake rate of M0 and M1 | 0 | $min^{-1}$ | Estimated |
| $U_{debris}$[DC,M0,CD8+] | debris uptake rate of DCs, M0, and CD8+ T cells | 0.1 | $min^{-1}$ | [1] |
| $S_{TNF}$[M1,M2] | TNF secretion rate of M0 and M1 | 1 | $min^{-1}$ | Estimated |
| $S_{debris}$[Dead cells] | debris secretion rate of all dead cells | 1 | $min^{-1}$ | [1] |
| $r_{recruit}$[MP] | recruitment rate of macrophages | $4.0 \times 10^{-9}$ | $cell/min \cdot \mu m^3$ | [1] |
| $r_{recruit}$[DC] | recruitment rate of dendritic cells | $2.0 \times 10^{-9}$ | $cell/min \cdot \mu m^3$ | [1] |
| $\rho_{min}$[MP] | minimal signal of TNF to recruit macrophages | 0.1 | TNF concentration | [1] |
| $\rho_{min}$[DC] | minimal signal of TNF to recruit DCs | 0.1 | TNF concentration | [1] |
| $\rho_{sat}$[MP] | saturation signal of TNF to recruit macrophages | 0.3 | TNF concentration | [1] |
| $\rho_{sat}$[DC] | saturation signal of TNF to recruit DCs | 0.3 | TNF concentration | [1] |

**Table S2**: Table of phagocytosis, attachment, mechanical parameters.

| Parameter | Description | Value | Unit | Reference |
|---|---|---|---|---|
| $r_{phag}$ | phagocytosis rate of macrophages | 0.167 | $min^{-1}$ | [1] |
| $V_{max}$ | Max volume of macrophages to stop cell engulfment | $6.5 \times 10^3$ | $\mu m^3$ | [1] |
| $r_{death}$ | death rate of exhausted macrophages | 0.01 | $min^{-1}$ | [1] |
| $r_{V_{int}}$ | intracellular digestion rate in macrophages | 1 | $\mu m^3/min$ | [1] |
| $r_{attach}$[DC] | attachment rate of dendritic cells | 100.0 | $min^{-1}$ | Estimated |
| $r_{leave}$ | DC exit rate from the tumor microenvironment | 0.2 | $min^{-1}$ | Estimated |
| $\bar{r}_{01}$[CD8+ nearby DC] | DC-induced CD8+ T cell cycle entry rate | $2.08 \times 10^{-3}$ | $min^{-1}$ | [1] |
| $r_{attach}$[CD8+ nearby DC] | attachment rate of CD8+ induced by DC | 0.6 | $min^{-1}$ | [1] |
| $r_{attach}$[CD8+] | attachment rate of CD8+ | 0.2 | $min^{-1}$ | [1] |
| $T_{attach}$ | the mean duration of CD8+ and cancer cells attachment | 8.5 | $min$ | [1] |
| $r_E$ | coefficient for the magnitude of the elastic force | $5.0 \times 10^{-2}$ | $min^{-1}$ | [2] |
| $r_P$ | relaxation rate according a plastic reorganization | $5.0 \times 10^{-4}$ | $min^{-1}$ | [2] |
| $d_P$ | maximum deformation of parenchymal cells | 0.75 | $\mu m$ | [2] |
| $\bar{r}_{01}$ | base rate for entry of cancer cells into the cell cycle | $2.0 \times 10^{-3}$ | $min^{-1}$ | Estimated |
| $p_{max}$ | maximum simple pressure to cancer cells proliferate | 10 | – | Estimated |

**Table S3**: Table of parameters associated with lymph node and tumor microenvironment interaction.

| Parameter | Description | Value | Unit | Reference |
|---|---|---|---|---|
| $\tau_{DC}$ | time to DCs migrate to lymph node | 720 | $min$ | [1] |
| $k_D$ | antigen presentation rate | 1 | $min^{-1}$ | [1] |
| $\delta_{DM}$ | natural decay rate of $D_M$ | $3.5 \times 10^{-4}$ | $min^{-1}$ | [1] |
| $\sigma_{T_{H1}}$ | proliferation rate of $T_{H1}$ | $7.0 \times 10^{-4}$ | $cells^2/min$ | [1] |
| $\pi_{T_{H1}}$ | activation rate of $T_{H1}$ | $1.5 \times 10^{-5}$ | $min^{-1}$ | [1] |
| $\delta_{T_{H1}}$ | deactivation rate of $T_{H1}$ | $7.0 \times 10^{-7}$ | $cell^{-1}min^{-1}$ | [1] |
| $\beta$ | half maximum deactivation of $T_{H1}$ | $5.0 \times 10^{2}$ | $cells$ | [1] |
| $\mu_{T_H}$ | natural death rate of $T_{H1}$ and $T_{H2}$ | $1.56 \times 10^{-5}$ | $min^{-1}$ | [1] |
| $\sigma_{T_{H2}}$ | proliferation rate of $T_{H2}$ | $2.8 \times 10^{-5}$ | $cells/min$ | [1] |
| $\pi_{T_{H2}}$ | activation rate of $T_{H2}$ | $2.0 \times 10^{-6}$ | $cell^{-1}min^{-1}$ | [1] |
| $\rho$ | weight conversion between $T_{H1}$ and $T_{H2}$ activation | 1 | $cell$ | [1] |
| $\pi_T$ | activation rate of T cells | $3.0 \times 10^{-3}$ | $min^{-1}$ | [1] |
| $\phi_T$ | scaling factor between lymph node and tumor microenvironment | $1.0 \times 10^{6}$ | $cells$ | [1] |
| $\beta_{T1}$ | half maximum of activation of T cells | $2.0 \times 10^{3}$ | $cells$ | [1] |
| $\delta_T$ | deactivation rate of T cells | $7.0 \times 10^{-4}$ | $min^{-1}$ | [1] |
| $\beta_{T_2}$ | half maximum deactivation of T cells | $2.0 \times 10^{2}$ | $cells$ | [1] |
| $\delta_C$ | natural decay rate of T cells | $1.4 \times 10^{-6}$ | $min^{-1}$ | [1] |
| $\phi_C$ | population threshold of T cells | 10 | cells | [1] |
| $\kappa_T$ | recruitment rate of $T_{Ct}$ and $T_{ht}$ | $1.1 \times 10^{-4}$ | $min^{-1}$ | [1] |
| $\tau_{TC}$ | time to $T_{Ct}$ and $T_{ht}$ infiltrate the tumor microenvironment | 360 | $min$ | [1] |

**Table S4**: Range of parameters of virtual patients.

| Parameter | Description | [Min, Max] | Unit |
|---|---|---|---|
| $r_{recruit}[MP]$ | recruitment rate of MP | $[0, 8.0 \times 10^{-9}]$ | $cell/min \cdot \mu m^3$ |
| $\rho_{min}[MP]$ | minimal signal of TNF to recruit MP | $[0, 0.2]$ | TNF conc. |
| $\rho_{sat}[MP]$ | saturation signal of TNF to recruit MP | $[0, 0.6]$ | TNF conc. |
| $r_{recruit}[DC]$ | recruitment rate of DCs | $[0, 4.0 \times 10^{-9}]$ | $cell/min \cdot \mu m^3$ |
| $\rho_{min}[DC]$ | minimal signal of TNF to recruit DCs | $[0, 0.2]$ | TNF conc. |
| $\rho_{sat}[DC]$ | saturation signal of TNF to recruit DCs | $[0, 0.6]$ | TNF conc. |
| $r_{leave}$ | DC exit rate from the TME | $[0, 0.4$ | $min^{-1}$ |
| $\delta_C$ | natural decay rate of T cells | $[0, 2.8 \times 10^{-6}]$ | $min^{-1}$ |
| $\kappa_T$ | recruitment rate of $T_{Ct}$ and $T_{ht}$ | $[0, 2.2 \times 10^{-4}]$ | $min^{-1}$ |
| $\delta_{DM}$ | natural decay rate of $D_M$ | $[0, 7.0 \times 10^{-4}]$ | $min^{-1}$ |

Subject to $\rho_{min}[DC] < \rho_{sat}[DC]$ and $\rho_{min}[MP] < \rho_{sat}[MP]$ constraints.

### 2 Data compression

The multiscale model generates high-dimensional data with spatial and temporal information. According to our specifications, each simulation stores 31 snapshot images over ten days of evolution, resulting in a total dataset size of approximately 5.55 TB. To simplify and reduce the dataset size, we perform a massive dimensionality reduction on the snapshot images, condensing the entire dataset to 6 GB. The arrangement of cells and substrates in the microenvironment at a specific simulated snapshot is encapsulated in a 24-bit 5x5 RGB bitmap image, where each channel encodes specific information:

- **Red channel (first 8 bits):** Stores information on the presence or absence of parenchymal cells, cancer cells, and dead immune cells.
- **Green channel (next 8 bits):** Represents the presence of live immune cells.
- **Blue channel (last 8 bits):** Specifies concentration ranges of TNF (tumor necrosis factor) and debris (as defined in Table S5).

For example, consider a pixel where there are live cancer cells, macrophage M0, and voxels with debris concentrations in the range of [0.2, 0.3], with a TNF concentration 0. According to Table S5, the pixel color in RGB will be:

$$\begin{bmatrix} 0 & 0 & 0 & 0 & 0 & 0 & 0 & 1 \\ 0 & 1 & 0 & 0 & 0 & 0 & 0 & 0 \\ 1 & 1 & 0 & 0 & 1 & 0 & 0 & 0 \end{bmatrix} \begin{bmatrix} 2^0 \\ 2^1 \\ \vdots \\ 2^7 \end{bmatrix} = \begin{bmatrix} 128 \\ 2 \\ 19 \end{bmatrix}$$

This approach allows us to drastically reduce the output space of the model, admitting it comes at the cost of losing some data resolution while retaining crucial aspects of the spatial and temporal dynamics.

The conversion process is accomplished by mapping the computational domain into partitions that define the pixels of the bitmap image. Within each pixel, we calculate the presence (bit 1) or absence (bit 0) of specific cell phenotypes and concentrations of relevant chemicals, such as TNF and debris. Figure S1 illustrates the data compression for three different immune control scenarios of micrometastases on day ten (NC, MC, and SC).

**Table S5**: Signature of the bits in RGB image during the process of data compression.

| Bit position | Red Channel | Green Channel | Blue Channel |
|---|---|---|---|
| 1 | live parechymal cell | inactivated DC | $0.00 \leq \rho_{debris} < 0.25$ |
| 2 | dead parechymal cell | macrophage M0 | $0.25 \leq \rho_{debris} < 0.50$ |
| 3 | dead cacner cell | exhausted MP | $0.50 \leq \rho_{debris} < 0.75$ |
| 4 | dead DC | hyperactivated MP | $\rho_{debris} \geq 0.75$ |
| 5 | dead MP | live CD4+ T cell | $0.00 \leq \rho_{TNF} < 0.25$ |
| 6 | dead CD4+ T cell | activated DC | $0.25 \leq \rho_{TNF} < 0.50$ |
| 7 | dead CD8+ T cell | activated MP | $0.50 \leq \rho_{TNF} < 0.75$ |
| 8 | live cancer cell | live CD8+ T cell | $\rho_{TNF} \geq 0.75$ |

DC: dendritic cell, M0: inactivated macrophage, MP: macrophage, $\rho_{TNF}$: concentration of tumor necrosis factor, $\rho_{debris}$: concentration of debris.

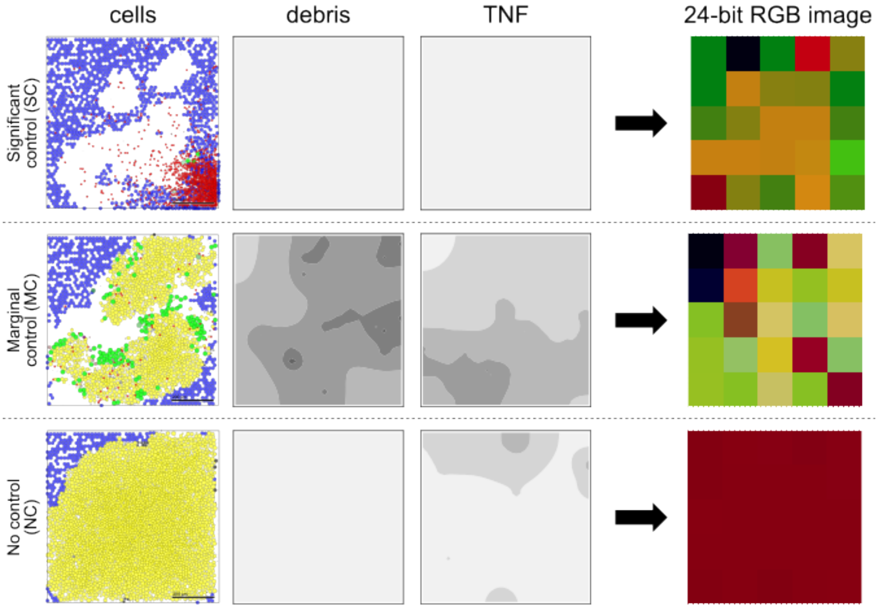


**Fig. S1**: The spatial data compression into a 24-bit RGB image from the multiscale model of immune surveillance in micrometastases.

### 3 Analysis of clustering of all trajectories

We conducted an investigation comparing two distinct clusters of NC-labeled trajectories resulting from dimensionality reduction using Uniform Manifold Approximation and Projection (UMAP) across a dataset of 100,000 virtual patient trajectories. Each trajectory is represented by 31 snapshots of 5x5 bitmap RGB images. Our findings reveal significant differences in immune system activation between these clusters. Cluster 1, located on the left side in Figure S2, exhibits a heightened level of immune system activation. In contrast, Cluster 2 on the right side in Figure S2 displays a decline in immune system activity. Our comprehensive analysis underscores differences in the composition of immune cell populations within these two samples. Upon obtaining additional samples, we discerned that the primary dissimilarity lies in the production of T cells within the lymph nodes. Cluster 1 consistently exhibits an upward trajectory in T cell production, while Cluster 2 reaches a state of saturation. T cell production relies entirely on immune activation, specifically driven by the activation and migration of dendritic cells (DCs). These DCs, in turn, trigger T cell production.

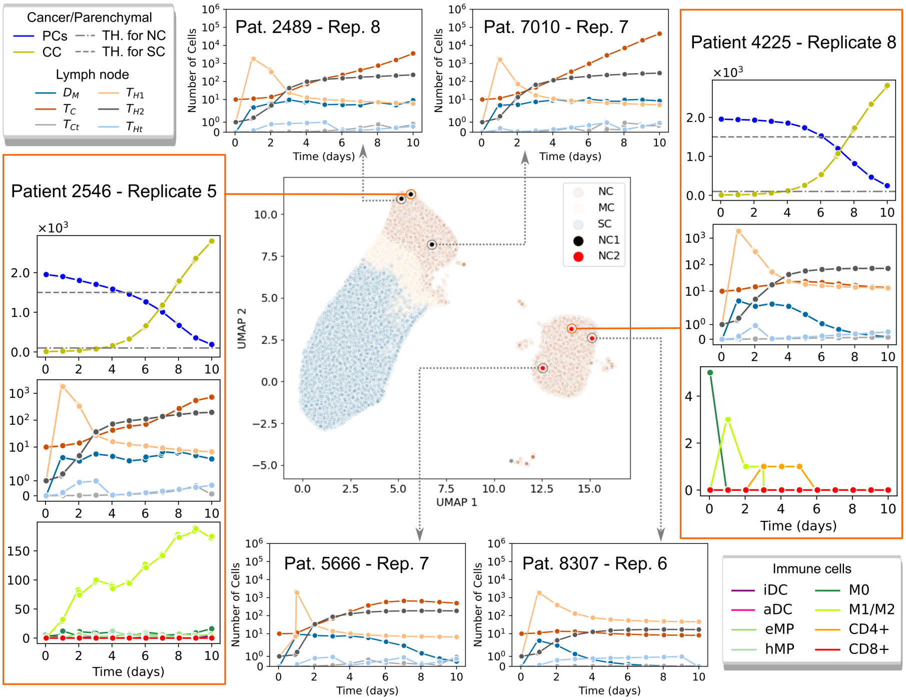


**Fig. S2**: Visualization of Spatial and Temporal Trajectories in the No Control (NC) Case. In the reduced space represented by Uniform Manifold Approximation and Projection (UMAP), this figure showcases the spatial/temporal trajectories of all patients in the no control (NC) case. The samples denoted by orange arrows correspond to cell population plots of cancer cells, parenchymal cells, lymph node (LN) cells, and immune cells within the microenvironment. Meanwhile, the samples marked with gray arrows emphasize distinctions between clusters. Notably, the cluster on the right in the NC case exhibits a higher population of T cells within the lymph node compared to the left cluster, indicative of immune system activation.

### 4 Classification analysis

We perform time series clustering with the Python library tslearn [3] with k-means and the dedicated time series metric, Dynamic Time Warping (DTW). This metric effectively addresses the issue of time shifts dependencies in the trajectories. In the confusion matrix (Table S6), we compare this clustering with the classifier strategy adopted in this work, based on a cancer population thresholds on day 10 of the simulation used in this manuscript. Notably, both classifications exhibit considerable similarities, with Cluster 0 and 1 capturing our extreme case labels significant control (SC) and no control (NC), respectively, with over 90% accuracy. Figure S3 corroborates this finding, displaying the average and standard deviation for each group along with the

100,000 trajectories. Altogether, this analysis suggests that our classifier have low dependence on individual time variations but relies on the entire trajectory dynamics.

**Table S6**: Normalized confusion matrix for the classification based on the threshold of the cancer cell population on day 10, along with the time series clustering over a 10-day period using DTW.

| | Sign. Control (SC) | No Control (NC) | Marg. Control (MC) | Total$^\star$ |
|---|---|---|---|---|
| Cluster 0 | 0.92 | 0.00 | 0.08 | 41,035 |
| Cluster 1 | 0.00 | 1.00 | 0.00 | 44,775 |
| Cluster 2 | 0.27 | 0.03 | 0.70 | 14,190 |
| Total$^\star$ | 41,808 | 45,149 | 13,043 | 100,000 |

$^\star$ Total number of trajectories.

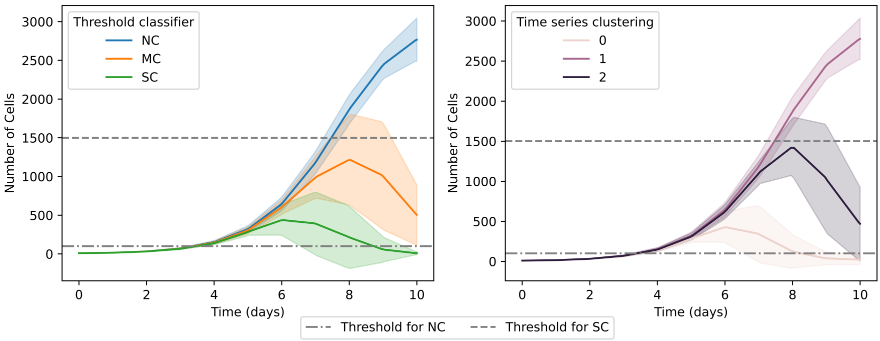


**Fig. S3**: The comparison of live cancer cell population over time. Data obtained from 100,000 trajectories using our patient classifier and dynamic time warping.

**Table S7**: Trajectories of four patients displaying different immune response classifications. Patient 6 shows multiple labels across trajectories (NC, SC, MC), patient 2 shows all trajectories classified as no control (NC), patient 93 shows all trajectories classified as marginal control (MC), and patient 5 shows all trajectories classified as significant control (SC). Each row is organized by the specific label.

| ID | Label | Cancer/Parenchymal | Lymph Node | Immune |
|---|---|---|---|---|
| 6 | NC | 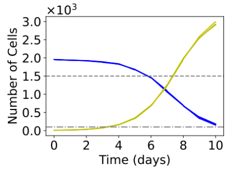 | 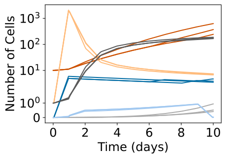 | 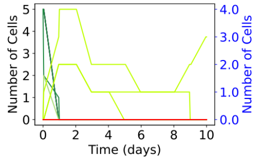 |
| 6 | MC | 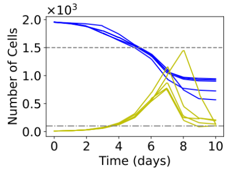 | 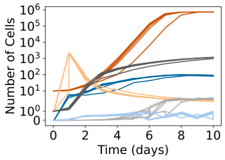 | 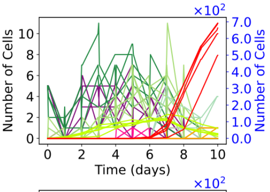 |
| 6 | SC | 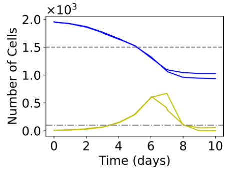 | 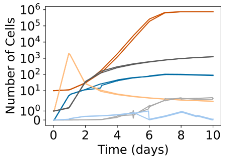 | 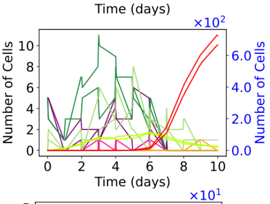 |
| 2 | NC | 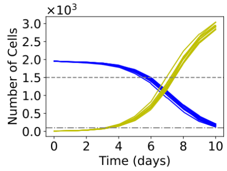 | 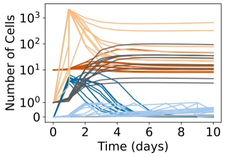 | 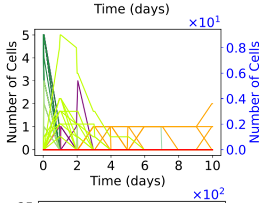 |
| 93 | MC | 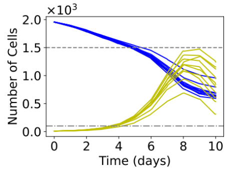 | 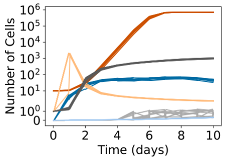 | 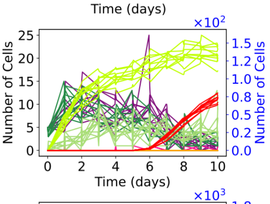 |
| 5 | SC | 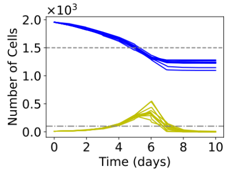 | 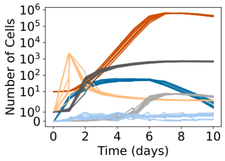 | 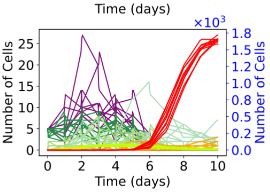 |

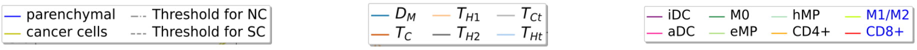


$\mathbf{D_M}$: DCs that have migrated from the TME to the LN, $\mathbf{T_{H1}}$: helper T cells type one, $\mathbf{T_{H2}}$: helper T cells type two, $\mathbf{T_C}$: cytotoxic T cells, $\mathbf{T_{Ct}}$: CD8+ cytotoxic T cells, $\mathbf{T_{ht}}$: CD4+ helper T cells, **iDC**: inactive DC, **aDC**: active DC, **eMP**: exhausted macrophages, and **hMP**: hyperactivated macrophages.
